# Plant-animal below-ground interaction modifies plant phenotype and its above-ground interaction: a review and new case study

**DOI:** 10.1101/2024.04.29.591669

**Authors:** Celia Vaca-Benito, Camilo Ferrón, Antonio J. Velázquez de Castro, A. Jesús Muñoz-Pajares, Mohamed Abdelaziz, Ana García-Muñoz

## Abstract

Ecological interactions play a role in promoting and maintaining biodiversity. These interactions form complex networks of interconnected species. Therefore, changes resulting from an interaction in one of the partners can have indirect consequences on subsequent interactions with other species. Since the mutualism-antagonism continuum is a gradient, a shift in the strength and sign of an interaction is possible, highlighting the dynamic nature of interaction networks. In flowering plants, a wide variety of below- and above-ground interactions are established with a single host plant. Changes in the host, derived from such interactions, can modulate the outcome of the remaining connections in both strength and sign, and the overall configuration of the network. Thus, a species can mediate community-wide consequences through its interaction with the host by altering the plant phenotype. We present a case study where a root infection has unexpected consequences on the pollination host, driving phenotypic changes. This study provides new data on the dynamism of species interactions and the importance of obtaining a global view of interaction networks. Disentangling the direct and indirect effects of interactions and their impact on the rest of the interactions in wild communities is essential for a good understanding of the evolutionary and ecological mechanisms that promote and maintain biodiversity.

## Introduction

Species diversification and ecological interactions have been intertwined throughout the evolutionary history of organisms (Thompson 1999). Species are interconnected, giving rise to a broad range of associations. In the wild, these associations are not isolated but instead form complex networks of interacting species, where each relationship between two organisms can simultaneously influence others (Pilosof et al. 2017). The direct and indirect effects of these interactions on individual fitness can function as selective pressures and even modify adaptive trajectories within a population or community. However, predicting the totality of ecological interactions and understanding how one interaction can modify or promote another is challenging.

The network of interactions in nature is diverse and multi-dimensional. Furthermore, the magnitude and direction of such interactions are dynamic, as each one is affected directly or indirectly by external factors (Sawaya et al. 2018; Moran et al. 2022). In other words, the outcomes of species interactions are context-dependent (Chamberlain et al. 2014; Song et al. 2020). For instance, global climate change impacts the configuration of species interactions (Tylianakis et al. 2008) because organisms respond plastically by altering their phenotypes (Stotz et al. 2021). Likewise, biotic factors can alter phenotypes and thus subsequent interactions (Andriuzzi et al. 2016). An interesting example is the interaction between the perennial herb *Ruellia nudiflora* (Engel. & Gray) and larvae of the *Tripudia* moth species, its main adversaries. However, the strength of this antagonistic interaction is mitigated due to parasitism of these larvae by wasps and fly species from different families (Moreira et al. 2015). In this sense, intraspecific phenotypic variation has been proposed as a mechanism capable of altering the links among different types of interactions (Moran et al. 2022). Since individuals, rather than species themselves, are the components of interactions (Jordano 2016a), intraspecific variation is particularly relevant in wild communities (Nakazawa 2017; Arroyo-Correa et al. 2023). Thus, identifying inter-individual variation and other mechanisms that underlie the direction and strength of ecological interactions is essential to understand the evolutionary implications of species relationships.

Despite extensive literature on species interactions, characterizations of ecological networks involving more than two partners or species at the level of fitness components are rare (Table 1). Our aim in this work is to explore the wide variety of species interactions briefly and highlight the limited knowledge on how indirect effects of one interaction can alter subsequent relationships with other species. We have focused mainly on flowering plants due to the extensive variety of associations that a host can establish both below and above ground. Finally, we present a case study illustrating unexpected effects of an antagonistic interaction affecting additional relationships and potentially the population structure and dynamics of an autogamous plant species.

**Table 1.** Summary of species interaction networks established in plants involving only two known partners (shadow cells) and more than two partners (white cells). Shadow cells represent only a brief proportion of the total known simple interactions, which are often well-characterized. In contrast, complex networks are rarer and less characterized, with only a few examples where the different effects are identified. Interestingly, pollinators are often involved in such interactions. The letter "H" denotes the host species, and the positive or negative sign between parentheses indicates mutualistic or antagonistic interaction, respectively.

| INTERACTING SPECIES | CONSEQUENCES | REFERENCES |
| --- | --- | --- |
| H: *Vigna unguolata*, *Lonchocarpus sericeus*,<br>*Sesbania rostrata*,<br>*Tephrosia platycarpa*.<br>(-) *Maruca vitrata*<br>(+) *Therophilus javanus* | Herbivory stimulates volatiles emission that attract wasp parasitoids of the herbivore larvae | Dannon *et al.* (2010)<br>Aboubakar Souna *et al.* (2019) |
| H: *Silene latifolia*<br>(-) *Spodoptera littoralis*<br>(+) nocturnal moths | Folivory changes floral signaling which attract more pollinators | Cozzolino *et al.* (2015) |
| H: *Brassica nigra*<br>(-) *Pieris brassicae*<br>(+) *Apis mellifera*, *Epishyrphus balteatus* | Folivory reduces the duration of pollinator visits | Bruinsma *et al.* (2014) |
| H: *Eryngium yuccifolium*<br>(-) *Coletechnites eryngiella*<br>(+) generalist pollinators | Herbivory reduces inflorescence size and pollinator visits | Danderson & Molano-Flores (2010) |
| H: *Centrosema virgianum*<br>(-) blister beetles and Agromyzidae flies<br>(+) *Bombus pensylvanicus*, *Xylocopa micans*,<br>*Megachile spp.*, *Melissodes spp.* | Folivory decreases the frequency of pollinator visits | Cardel & Koptur (2010) |
| H: *Macaranga tanarius*<br>(-) herbivores species<br>(+) ant and wasp species | Extrafloral nectar nourishes ants that defend their host from herbivores | Heil *et al.* (2000, 2001) |
| H: *Asclepias spp.*<br>(-) *Aphis nerii*, *Danaus plexippus*<br>(+) arbuscular mycorrhizal fungi (AMF) | AMF activity produces toxic leaves against herbivory which also affect root AMF | Meier & Hunter (2018) |
| H: *Quercus ilex*, *Q. faginea*.<br>(+) *Tuber melanosporum* | Improvement of seedling growth and water and phosphorus uptake | Núñez *et al.* (2006) |
| H: *A. thaliana*, *Populus tremula*, *P. alba*<br>(+) *Laccaria bicolor* | Development of lateral roots and improvement of resources usage | Felten *et al.* (2009) |
| H: *Asclepias spp.*<br>(-) AMF | Improvement of plant defense against herbivores | Vannette *et al.* (2013) |
| H: *Coffea arabica*, *C. canephora*<br>(-) *Hemileia vastatrix* | Premature leaves fall and photosynthesis decrease | Silva *et al.* (2006) |
| H: *Solanum lycopersicum*<br>(-) Cucumber mosaic virus (CMV) | Malformation and fructification decrease | Morales *et al.* (2009) |
| H: *Arabidopsis thaliana*<br>(+) *Pseudomonas fluorescens* | Improvement of plant growth and disease control | Lugtenberg & Kamilova (2009) |

### How do species interactions occur?

Species interactions range from mutualism to antagonism, although more neutral associations and intermediate values also exist along this gradient (Gómez et al. 2023). Mutualism involves reciprocally positive interactions between pairs of species (Bronstein 2009; Bronstein 2015). Animal-mediated pollination is one of the most well-known examples of mutualism in flowering plants (Herrera and Pellmyr 2009). Since pollinator activity is influenced by preferences for specific floral structures, plant intraspecific variation serves as a substrate for natural selection driven by flower visitors (Grant 1949; Muchhala and Potts 2007; Rymer et al. 2010; Van der Niet et al. 2014; Minnaar et al. 2019). In contrast, antagonistic interactions such as parasitism or herbivory have adverse effects on the growth, reproduction, survival, or overall fitness of one of the organisms involved (Preston 2010). Yet, trade-offs among beneficial and harmful organisms associated with roots have also been described (Detrey et al. 2022). The intensity and direction of selection are conditioned by the interplay between ecological and intrinsic factors, such as the phenotypes of both species (Fordyce, 2006). Thus, any interaction along the gradient (from mutualistic to antagonistic interactions) can become an important driver of species processes since they constitute strong selective pressures (Price et al. 1986).

Consequences of the same interactions can also vary according to context. In some palm populations, herbivory by moths had a positive effect on flowering and pollination, while herbivory by goats had the opposite effect (Muñoz-Gallego et al. 2022). This system illustrates how the same type of interaction exerted by two different species can have contrasting results on the host, at least in this population. Additionally, since fitness has several components, the differential effect of each herbivore species on each of these components can result in a net positive balance on fitness (Muñoz-Gallego et al. 2022). This balance between antagonistic relationships and host reproduction allows for the maintenance of such interactions, which is essential in highly specialized organisms. Other species exhibit strictly mutualistic interactions with a high degree of specialization. For example, *Frankliniella intonsa* (Trybom, 1895) thrips pollinate *Stellera chamaejasme* L. plants where they also find brooding sites and food for their offspring. This close relationship drives a coevolution process that fits the life cycles of both species (Zhang et al. 2021).

Ecological interactions also vary between specialized and generalist ones. This characterization attends to the number of links with diverse partners established by a species (Blüthgen et al. 2006). The level of specialization has significant consequences on the selective pressures resulting from these interactions (Ali and Agrawal, 2012), having the ability to modify the architecture of community assembly (Ponisio et al., 2019; Yeakel et al., 2020). Highly specialized species systems play an important role in maintaining species interactions and enhancing diversification patterns (Manzaneda and Rey 2009). Therefore, the degree of specialization of mutualistic and antagonistic interactions plays an important role in species coevolution (Ali and Agrawal 2012; Waser and Ollerton 2006). Within each type of relationship, obligatory and facultative interactions are also differentiated according to species dependency (Poisot et al. 2013). Moreover, intra-individual shifts in species interactions are expected across seasons, throughout the host life cycle in pollinator-plant systems (Valverde et al. 2016) or during seed dispersal processes (Wang and Smith 2002; Elzinga et al. 2007). Likewise, the ultimate effects of other types of interactions such as any kind of herbivory (e.g., florivory, frugivory, granivory, etc.) are context-dependent and thus subjected to spatio-temporal variation (Perea et al. 2013) (Figure 1).

**Figure 1.**
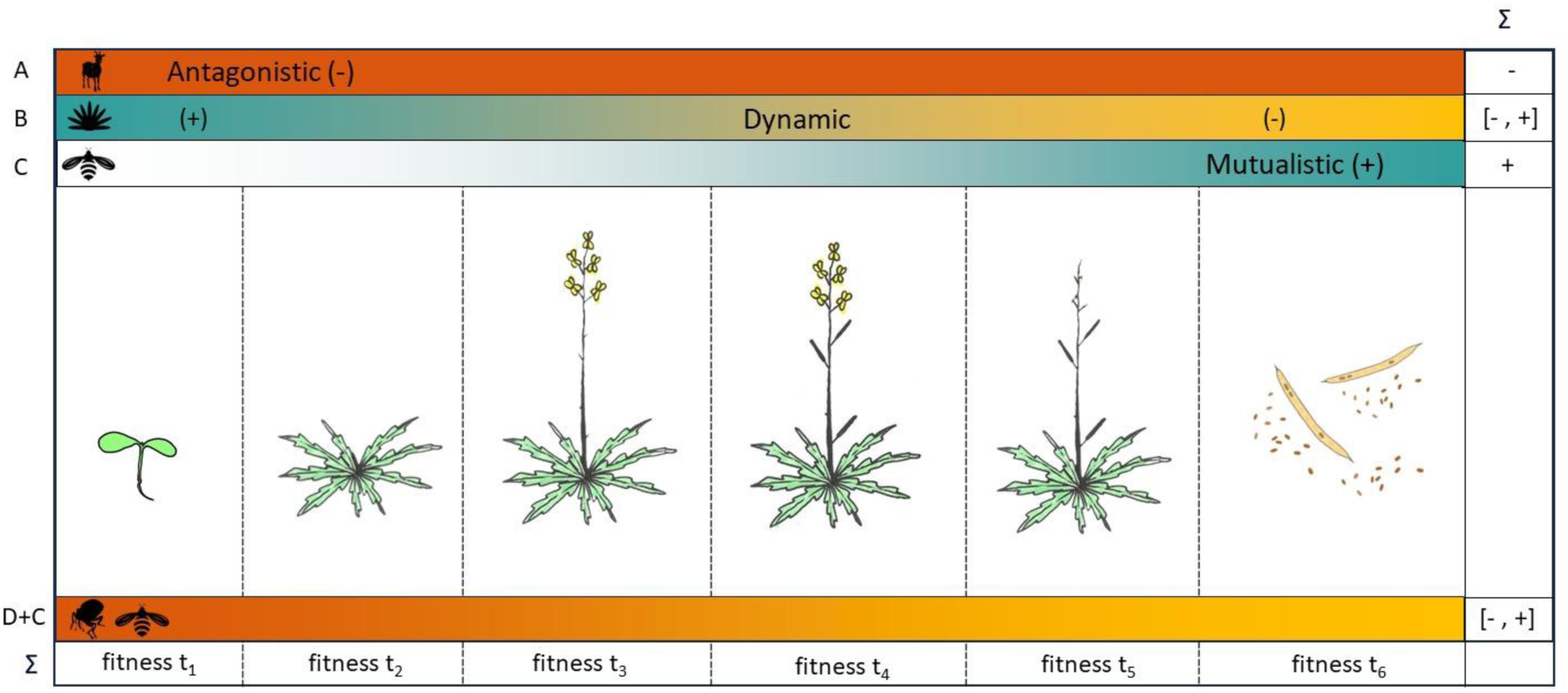
Representation of how the different stages of a host individual are subjected to changing effects of interactions with other species. *Fitness t_x_* indicates the fitness components of the plant (germination, growth, flowering, fruit development, and reproductive output as fruit and seed production). The summation (Σ) refers to the net effect of all interactions that can affect an individual throughout its entire life. Some examples of these dynamic interactions are: (A) Antagonistic, marked during the growth stage of the plant; (B) Dynamic interactions that can change from positive to negative effects on the hosts; (C) Mutualistic through insect-pollination during flowering; (D+C) When a species has a negative effect in earlier stages of the host’s life but promotes the positive effects of interactions established later with other species.

### Below-ground interactions

Individuals are exposed to a wide variety of interactions that simultaneously affect different organs. Often, these interactions exhibit opposite directions. Importantly, a significant portion of a plant’s body extends underground, forming roots. Therefore, many potential interactions established by a plant are barely visible to the naked eye, occurring below ground, where the complex network of primary and secondary roots interacts closely with soil biota. However, identifying the totality of soil organisms and their effects on roots remains a challenging issue, leading to misconceptions about many potential interactions. Nevertheless, extensive research on mycorrhizal fungi and rhizobacteria provides examples of mutualistic relationships between roots and soil organisms (Sapp 2004; Schulz 2006; Lugtenberg and Kamilova 2009; Mandyam and Jumpponen 2014; Rasmann and Turlings 2016). In contrast, detriments such as infections by parasites (Sukno et al. 2008; Quilbé et al. 2022) and herbivore attacks also occur in the root and provoke the activation of defense mechanisms (Lu et al. 2015).

An example of a specialist antagonistic interaction is herbivorous insects. Some are highly adapted to resist the secondary compounds of the plants they feed on. Some types of specialist herbivores can even manipulate their hosts. A notable example is gallers, which can reprogram the metabolism of their hosts to their advantage (Ali and Agrawal 2012). However, this reprogramming can have indirect effects on host phenotype, generating new selective opportunities and adaptive scenarios. Effectors that modulate plant defense responses have been identified in the saliva of gallers and other chewing herbivores (Hogenhout and Bos 2011). In general, root damage stimulates defense signaling responses that are different from those activated in above-ground host organs (Acosta et al. 2013; Johnson et al. 2016). Recent studies have documented the secretion of abscisic acid by larvae of gall-inducing insects for host plant manipulation, as this compound has significant consequences on plant growth and survival (Seng et al. 2023). Specifically, plants respond to root damage mainly by producing jasmonate (Lu et al. 2015), a plant hormone that triggers physiological cascades that can indirectly modulate other plant interactions, in addition to providing resistance against plant attackers (Toby Kiers et al. 2010; Lu et al. 2015).

### From below to above ground

Plants interact with a wide range of different organisms, collectively known as the phytobiome (Leach et al. 2017). The functionality of plant individuals strongly depends on roots due to their role in resource uptake and growth. Since most of the organisms that plants interact with live underground, any factor affecting root performance, such as parasitism and herbivory events, is expected to affect the above-ground plant body (Rasmann et al. 2017). In fact, both beneficial and detrimental organisms affecting roots are easily detectable through observation of the above-ground vegetative body. Additionally, species interactions established below ground can influence associations occurring above ground (Vannette and Hunter 2009). Mycorrhizal fungi are known to inhabit 90% of all plant species (Bonfante and Genre 2010), improving nutrient uptake (Meier and Hunter 2018). Furthermore, mycorrhizal fungi act as mediators in plant defense by modifying the expression of genes involved in physical structures and metabolic compounds of defense, dissuading herbivores and parasitoids even when they are specialists (Vannette and Hunter 2009, 2013; Vannette et al. 2013; Thamer et al. 2011). Despite the fact that interactions between below- and above-ground organisms occur, most studies on species interactions exclusively focus on above-ground interactions, overlooking the potential effects of the extensive below-ground biota (Van der Putten et al. 2001). Including the below-ground dimension in plant interaction studies would lead to a more realistic understanding of plant ecology, including those related to above-ground interactions.

### From benefits to detriments and vice versa

Unexpected consequences for overall performance are driven by the wide variety of associations established below and above ground. The dynamic nature of species interactions often falls within the ‘mutualism-parasitism-continuum’ (Saikkonen et al. 1998; Bronstein 2015), exposing a range in which shifts between antagonism and mutualism in both directions are possible. Phenotypic changes produced in host species by a given relationship influence the behavior of other interacting species and thus also the consequences of ongoing and subsequent associations. In flowering plants, the attack of herbivores or any pathogen on floral traits likely affects overall plant attractiveness (Moreira et al. 2019). As mentioned above, pollinators shape mutualistic relationships with flowering plants and ensure their reproduction. However, alterations in floral traits are expected to change pollinator preferences, decreasing visitation rates and undermining reproduction. Nevertheless, reproductive success is not always negatively affected by antagonists (Moreira et al. 2019). In fact, Cozzolino et al. (2015) found a good example of increased fitness as a secondary consequence of herbivore damage in *Silene latifolia*. Herbivory stimulates the emission of volatiles, which increases pollinator visitation in this system (Cozzolino et al. 2015), exemplifying a shift from a merely antagonistic interaction toward a beneficial one.

However, phenotypic changes in the host are not always directly caused by the attack, but a variety of defense mechanisms developed in harmed plants also affect host phenotype (Tiffin 2000). Phenotype can be directly affected by the release of phytohormones resulting from metabolic pathways activated in response to damage (Toby Kiers et al. 2010; Kammerhofer et al. 2015). Additionally, host plants exhibit compensatory mechanisms that reallocate resources to vegetative growth or reproduction (McNaughton 1983). Beyond their direct implications in defense and damage mitigation, these mechanisms can potentially have subsequent consequences on interactions with other species. For instance, chemical signals triggered during the attack directly deter pathogens and/or attract their natural predators via floral volatiles (Ekesi 2000; Dannon et al. 2010). However, costs derived from defense mechanisms have also been described (Altshuler 1999; Strauss et al. 1999; Ness 2006). This trade-off is expected when the mentioned compensatory mechanisms act by reallocating resources from reproductive investment to other functions (Strauss et al. 2002). Experiments in plant species tended by bodyguard ants also exemplify the conflicts that emerge from defense mechanisms involving more complex ecological networks. In these multispecies systems, ants exhibit defensive behavior against the herbivores of their host plant, *Ferocactus wislizeni* (Britton & Rose). However, ants also have a deterrent effect on their pollinators, affecting host reproduction (Ness 2006).

Since traits can be grouped by function, changes would have different implications according to the main traits involved in an interaction. As one might expect, below-ground traits such as roots are crucial for resource uptake, while vegetative traits are primarily affected by herbivory, and floral traits play a role in pollination. Nonetheless, the entire organism is more than the sum of its parts, which are closely interconnected. Unfortunately, following the potential implications of the variation caused by all interactions in a wild community is a challenging task. It becomes even more unfeasible when the complex metabolic networks underlying species interactions are unknown. Nonetheless, many researchers have outlined such complex interaction networks in plants. For example, although herbivory is expected to negatively affect the vegetative plant body, such as stems and leaves, it has been demonstrated to also influence the composition and amount of nectar, thus modulating pollinator behavior (Poveda et al. 2003; Gegear et al. 2007; Bruinsma et al. 2014). Changes produced in host individuals by interactions could have implications for the evolutionary trajectory of the species due to the presence of evolutionary dynamics in species interaction networks. Indeed, the parasite-driven wedge model was proposed as a mechanism driving parapatric speciation in animals (Thornhill and Fincher 2013). In animals, the behavior developed by host species to manage infection can function, in some cases, as a reproductive barrier. This is because assortative mating is controlled by behavior patterns, so gene flow can be limited among infected and non-infected individuals that differ in mating preferences.

### Phenotypic variation drives changes in species interactions

Inter-individual phenotypic variation is ubiquitous in any population. Such variation shapes the evolutionary potential as differences among individuals form the basis for natural selection to act upon and shape the evolutionary potential of the population (Caballero, 2020; Opedal et al., 2023). Thus, the selective pressure exerted by an organism with which the population interacts can be modulated by intra-individual variation. In this sense, the emergence of new phenotypes or changes in the frequency of existing ones could significantly alter the interactions that plants establish with the rest of the organisms in the community.

Phenotypic plasticity plays an important role in maintaining inter-individual variation within populations (Agrawal 2001). Besides abiotic factors, species interactions involve biotic aspects that can also affect phenotypic plasticity. For instance, plant individuals associated with endophytes show higher growth ratios and bigger flowers because of greater amounts of ethylene compared to plants without endophytes (Hardoim et al., 2008). Differences among individuals can drive a preference by the other partner of the association. In this way, phenotypic plasticity modulated by species interactions could also be enhanced by the inter-individual variation that affects the host choice by the other species.

### A new case of study

#### Study system

The autogamous species *Erysimum repandum* L., which belongs to the Brassicaceae family, is widely distributed across Europe and North America and associated with cereal crops (GBIF; www.gbif.org). This monocarpic plant shows small yellow flowers in which cross-pollination mediated by insects is not expected based on its selfing syndrome characteristics (Feliner 1991). Despite the small flowers expected in *E. repandum*, we observed inter-individual variation in flower and plant size in a wild population located in Serranía de Cuenca (Castilla-La Mancha, Spain; Figure 2A). In addition, a considerable number of individuals from this population presented a protuberance in the upper part of their axonomorphic main root (Figure 2B). This anomaly is the result of an infection by weevils (Coleoptera: Curculionoidea) whose larvae grow and feed (Figure 2C) within the root producing galls (Nieves-Aldrey, 1998). Weevils comprise the superfamily of beetles with the greatest richness of phytophagous species (Oberprieler et al. 2007), harming crops worldwide (Talekar 1982; Lu et al. 2015; Vale et al. 2023) although pollination activity of these animals has also been described (Haran et al. 2023).

**Figure 2.**
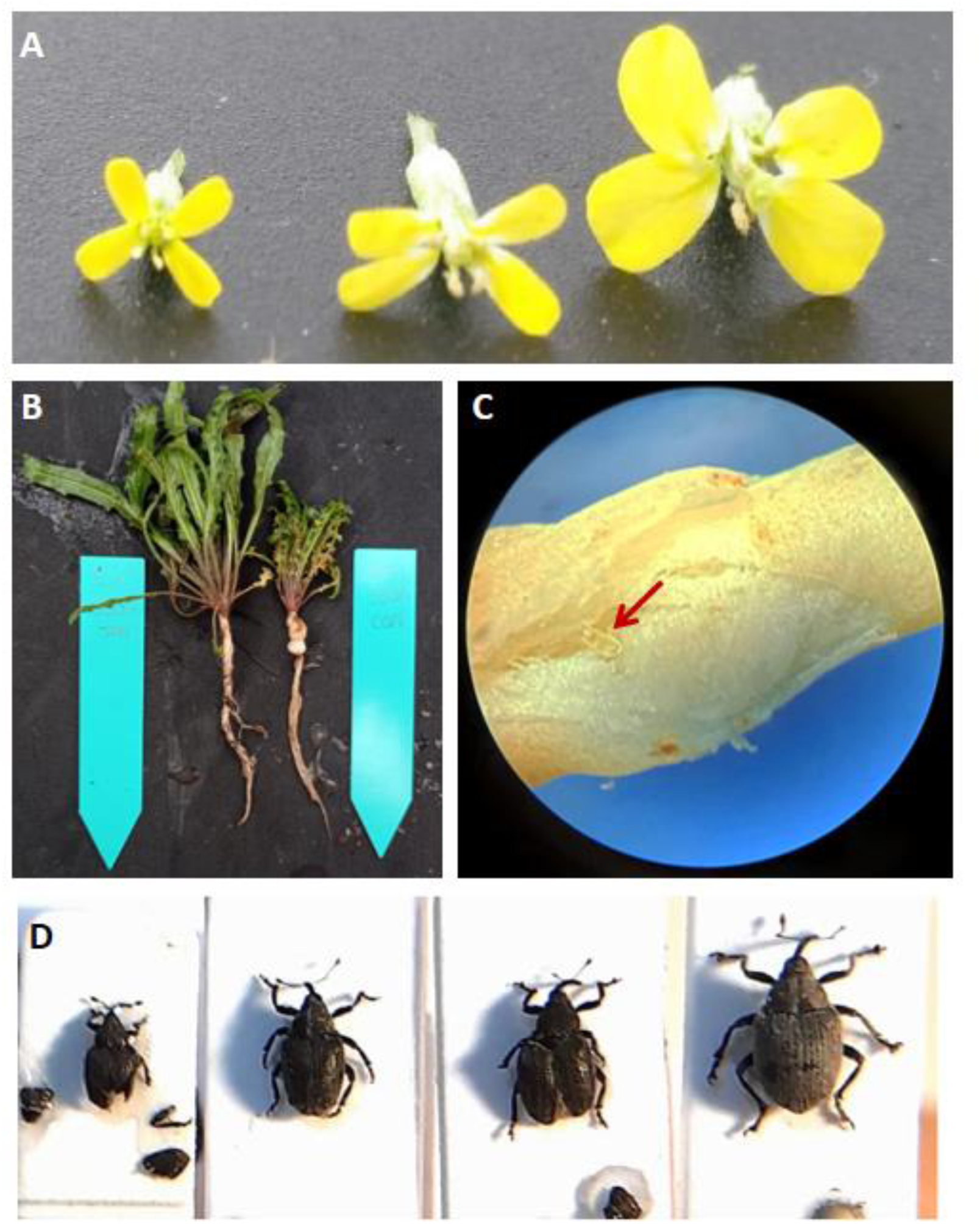
(A) Phenotypic variation in flower among different individuals from the same population of *E. repandum.* (B) Comparison between non-infected and infected root. (C) Visualization of an early-stage larva inside a galled root. (D) Collected individuals of *C. assimilis* (female and male), *C. picitarsis* and *C. napis*, respectively.

The gall presence in plants is expected to be detrimental for the host due to the larvae herbivory and the potential interference of the gall in the resources uptaken. Besides the formation of such structures, gall-inducing insects’ activity has molecular implications by releasing effectors which stimulate or inhibit phytohormones involved in plant growth and defense mechanisms (Giron et al. 2016). However, disentangling the reconfiguration of hormone networks induced by insect elicitors and predicting global effects on the host is not easy.

## Methods

We randomly marked and phenotyped individuals from our experimental population for two years (a total of 50 plants in 2022 and 90 plants in 2023). We mainly focused on (1) floral size, measured as petal and corolla length using a digital caliper (±0.01 mm error); (2) rewards production, measured as the nectar amount collected using glass Drummond® capillary 0.5 μL, and (3) plant size, as the height of the vegetative body measured using a flexometer (±0.5 cm error). In addition, we performed a pollinator census in 2022. We quantified the number of pollinator visits in each marked plant individual for 10-minute intervals. After taking all measurements, individuals were classified as infected and non-infected according to the presence or absence of root gall, respectively. Since gall root observation implied manipulating the individual, classification was the last step to avoid manipulation effects on measurements and pollinator visitation. Therefore, data imbalance between infected and non-infected plants was unavoidable.

To identify the weevil genus, we extracted DNA from galls in 2022 using the GenElute™ Plant Genomic DNA Kit (SIGMA) following manufacturer instructions. The purity, integrity, and concentration of the resulting DNA were evaluated by means of agarose gel visualizations and spectrophotometric methods (Qubit). Illumina libraries were performed using the Collibri ES DNA Library Prep Kit (ThermoFisher) and sequenced using a NovaSeq sequencing system. The sequences were aligned using Blastn filtering by Curculionidae in the GenBank database. In addition, weevil individuals were collected from the wild population in 2023 for the posterior taxonomic identification of the species.

Statistical analyses were conducted using ANOVA and Pearson’s product-moment correlation implemented in the stats package in R version 4.0.3 (R Core Team, 2023).

## Results

### Identification of interacting species

Genetic analyses revealed that the DNA found in galls corresponds to the *Ceutorhynchus* genus (Coleoptera). Due to the genome mix from galled roots, we could also identify the host plant and ensure that it was, in fact, *E. repandum* despite the observed size changes.

A morphologic study revealed a total of three different *Ceutorrhynchus* species from the isolated weevils in plants: *C. assimilis*, *C. picitarsis* and *C. napi* (Figure 2D). These three species are distributed in the western Palaearctic region. They are oligophagous on Brassicaceae (Dieckmann, 1972). Their larvae live inside the body of the plant and moves to the soil to pupate inside a cocoon (Scherf, 1964). *C. assimilis* (Paykull, 1792) is a small weevil (2.3 - 3.1 mm) whose body is entirely black or dark grey with scattered long pale scales. Its larvae produce big galls between stem and roots. (This species is widely referred to as *C. pleurostigma* [Marsham, 1802] in old works). *C. picitarsis* (Gyllenhal, 1837) is a little bigger (2.4 – 3.4 mm), adults may be recognized by the general coloration; entirely black with red tarsi. Eggs are laid on the underside of leaves at the bases of host petioles or on the stems, the freshly-emerged larvae feed on leaves for a while before they bore into soft stem tissue, here they feed throughout the winter, working their way down the stems as they do so, and end up in the root-collar when fully grown. *C. napi* (Gyllenhal, 1837) recognized by the big size (3.5 – 4.2 mm). Their larvae feed on the stem of the plant forming galls.

### Effects of the infection on plant morphology

Infected plants showed an anomaly in the root that could be hampering resource uptake, thus affecting plant health. However, contrary to expectations of a damaged appearance, infected plants exhibited an overall increase in plant size, measured as the height of the main flowering stalk, in both sampling years (Fig. 3).

**Figure 3.**
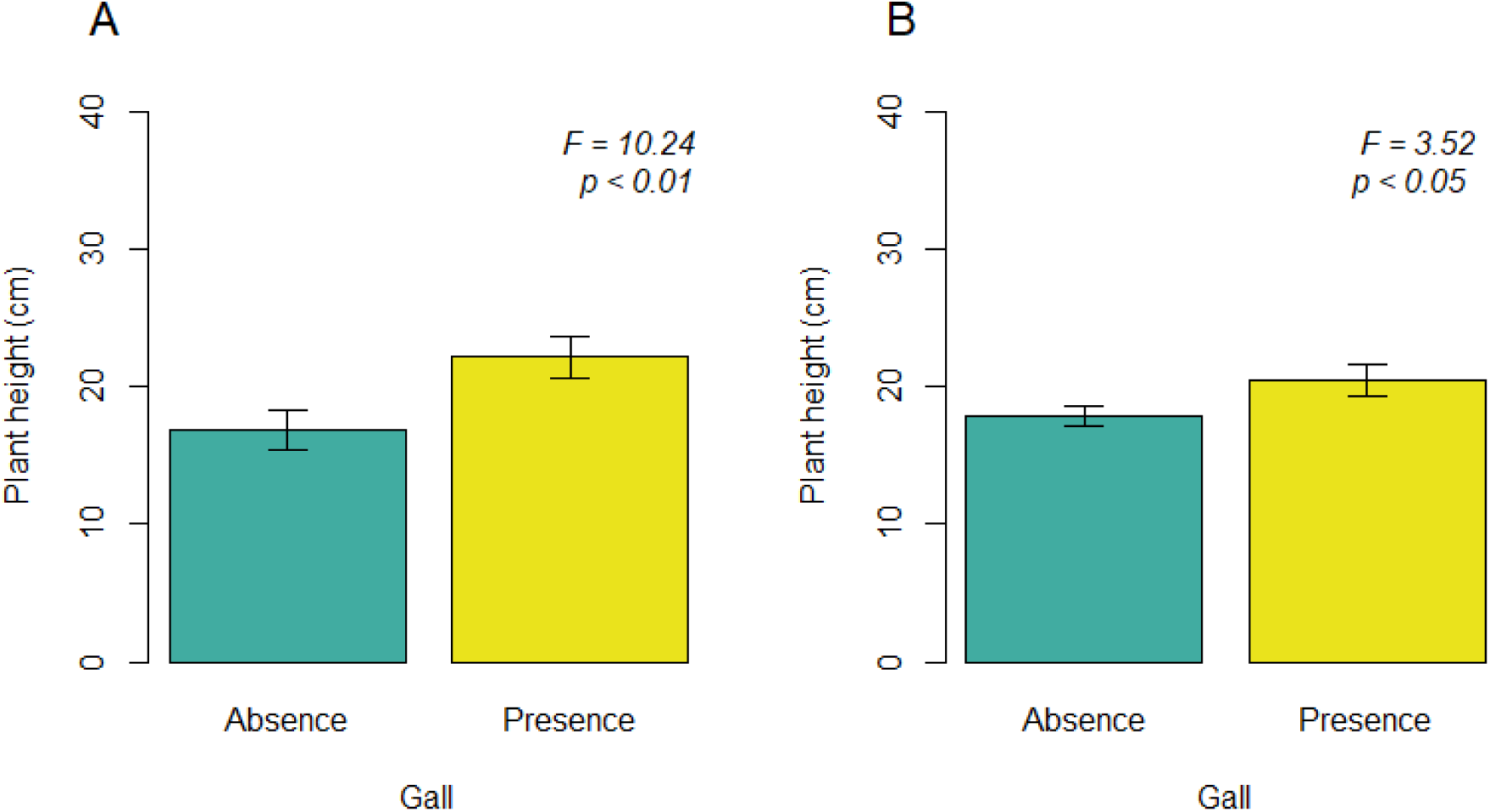
Differences in plant size, measured as height, between infected and non-infected plants for (A) the first sampling year in 2022 and (B) the second sampling year in 2023.

In addition to differences in the vegetative body of plants, we also observed variations at the floral level between individuals. Infected plants had significantly larger flowers than non-infected individuals in both the 2022 and 2023 sampling years (Fig. 4).

**Figure 4.**
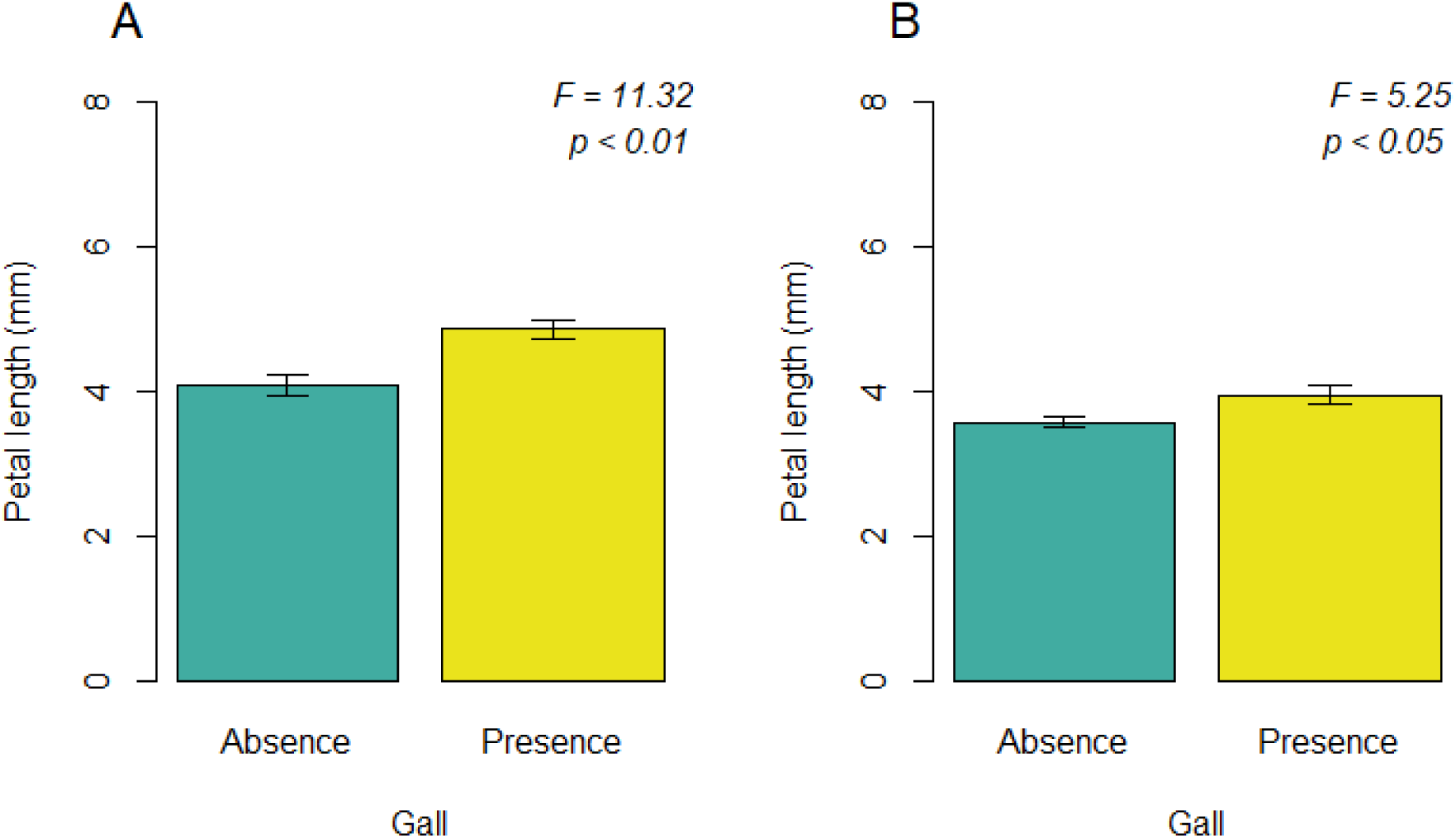
Differences in floral size, measured as petal length, between infected and non-infected plants for (A) the first sampling year in 2022 and (B) the second sampling year in 2023.

Inter-individual variation in both plant and flower size was easily observable. However, we also found significant differences in nectar production among individuals. Infected plants had flowers that produced more nectar than flowers from non-infected individuals (Fig. 5). This difference was maintenance for both years.

**Figure 5.**
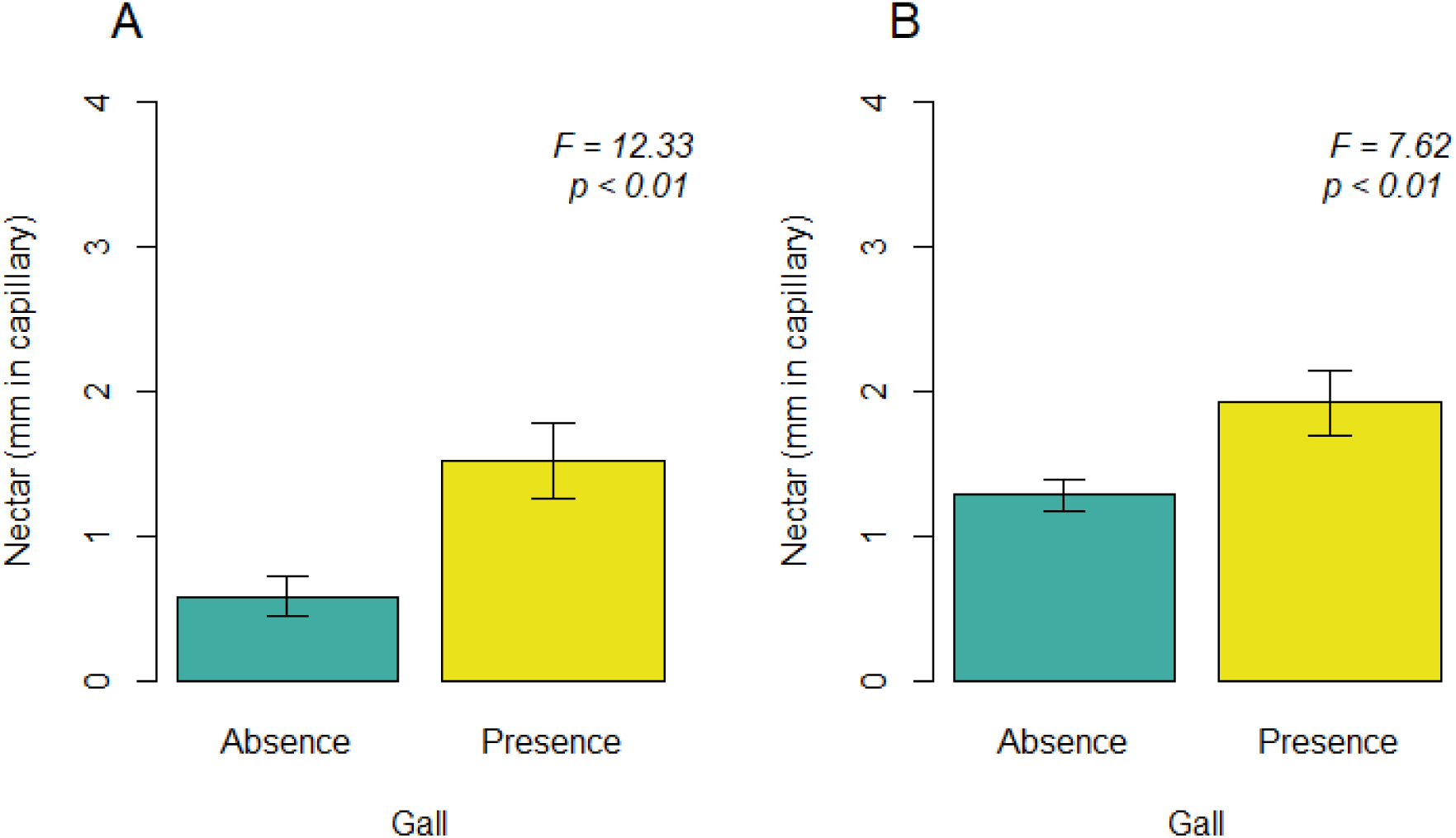
Differences in nectar amount per flower infected and non-infected plants for (A) the first sampling year in 2022 and (B) the second sampling year in 2023.

### Effect of phenotypic variation on pollinator interactions

Pollinators were not taxonomically classified, but the morphospecies were generally grouped as small bees, beetles, flies, and butterflies. While non-infected plants were scarcely visited, we observed a significantly higher preference by pollinators for infected plants (Fig. 6).

**Figure 6.**
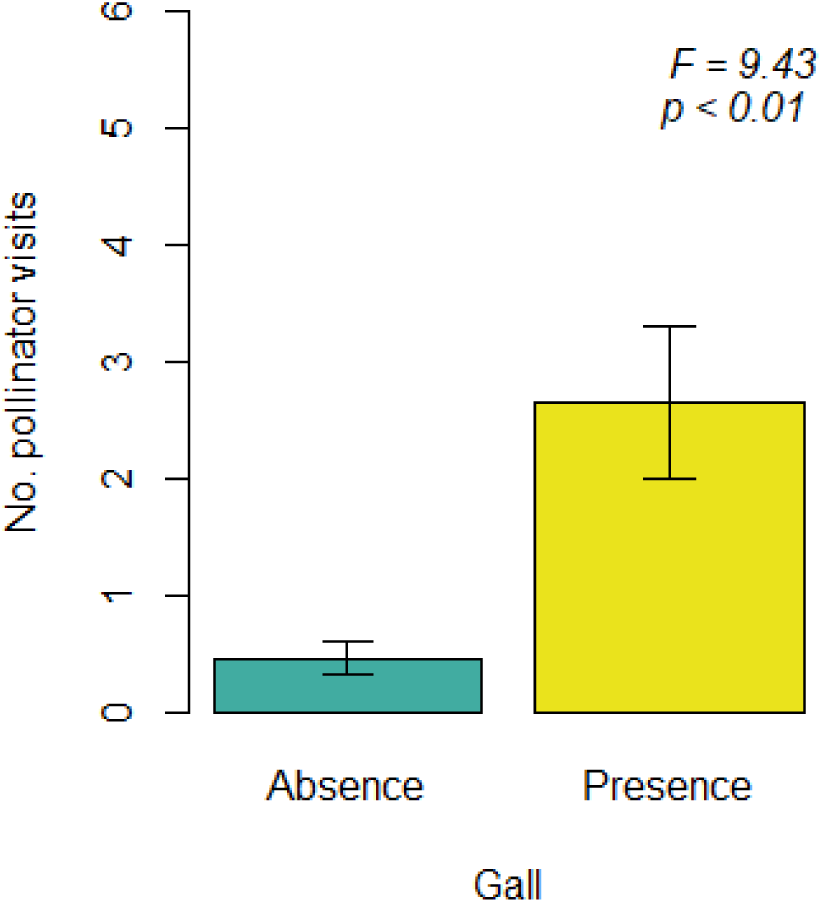
Differences in pollinator visits between infected and non-infected plants., measured as the number of insects that effectively touched the reproductive organs of flowers. Pollinators observation was performed only in the first sampling (2022).

In addition, we found that pollinators showed a marked preference for visiting individuals with larger flowers (Fig. 7A; t = 2.937, p-value < 0.01) and greater nectar amounts (Fig. 7B; t = 4.900, p-value < 0.0001). High values of these traits were exhibited by infected plants, indicating that an increase in floral and nectar traits, potentially influenced by infection, could affect pollinator behavior. Indeed, larger flowers seemed to be associated with greater nectar production (Fig. 7C).

**Figure 7.**
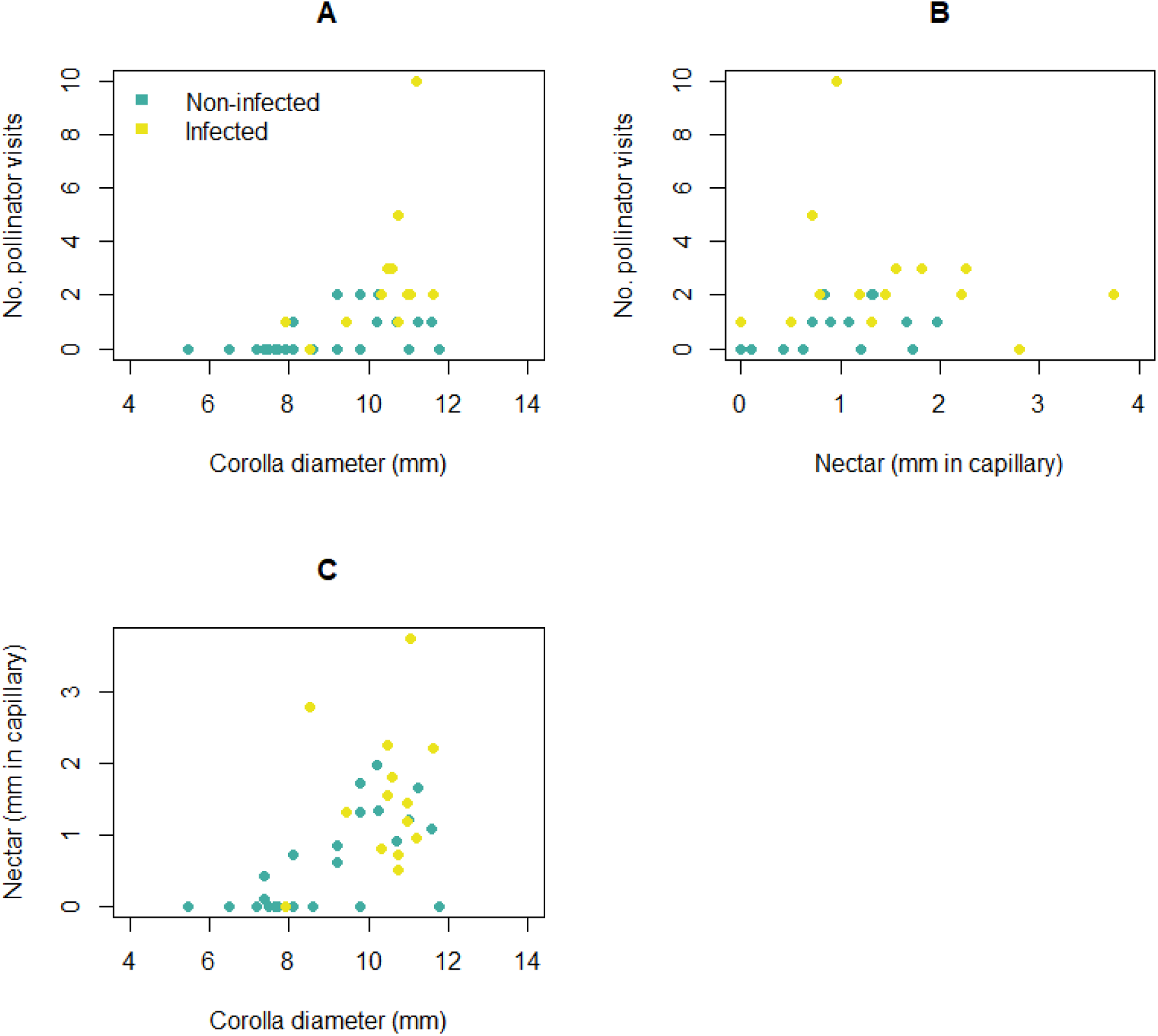
Correlation between (A) amount of pollinator visits and flower size measured as corolla diameter and (B) nectar production, and correlation between (C) nectar production and flower size.

## Discussion

The observed increase in flower size and reward is expected to potentially modulate the pollination ecology of infected plants. We recorded pollinator preferences for plants with larger flowers and greater nectar abundance within a population where both phenotypes coexist sympatrically. Pollination events are particularly notable in this population because *E. repandum* has traditionally been described as a self-pollinating species with no known pollinators. In fact, the authors of this work characterized the selfing syndrome of other *E. repandum* populations in the Iberian Peninsula and observed the absence of flower visitors during different samplings from 2015 to 2019 (authors’ personal observations).

Pollinator behavior would have consequences on population genetic structures due to gene flow among pollinated individuals. Given the expected lack of cross-pollination according to the original reproductive strategy of *E. repandum*, pollination events between infected plants would enhance outbreeding within the population. However, knowledge of the success of outcrossing events in *E. repandum* is needed to assess the extent of such genetic exchange. Although one might expect outbreeding to overcome the disadvantages of self-pollination, such as inbreeding depression, and increase genetic diversity, the opposite outcome might also be possible because the selfing evolutionary history of *E. repandum* would have ensured the purging of deleterious alleles. Nevertheless, evidence of foreign pollen tolerance and outcrossing success in other selfing *Erysimum* species has been documented. In any case, pollen movements between some individuals of the population might have important consequences for the mating system evolution of this species.

Far from appearing damaged, infected plants of *E. repandum* tended to have a greater total vegetative body size. An increase in vegetative tissue is expected as a response to herbivory through compensatory mechanisms. However, these mechanisms are primarily described when herbivory occurs in the vegetative parts rather than the roots. We observed a larger size of infected plants, which could compensate for the reduction in resource uptake by galled roots by increasing the photosynthetic fraction. However, an increase in corolla size and nectar abundance is not easily explained as a mechanism to counterbalance the loss of root functionality. Yet, changes in flowers could be explained as a consequence of the overall increase in plant size due to phenotypic correlation. Regarding nectar amount, it is noted that floral and nectar traits are often closely correlated, so genetic mechanisms such as pleiotropy and linkage disequilibrium could contribute to the indirect selection for other traits, such as nectar amount.

Since we found a strong correlation between root infection and phenotypic changes, we hypothesize that weevil infection is causing such phenotypes. However, the potential mechanisms underlying these associations are not yet identified, and thus, alternative explanations are plausible. Instead, intra-population variation could have a different origin, and weevils could simply be choosing larger plants or flowers to reproduce. To elucidate which mechanism is causing the observed changes, it is necessary to grow progeny of infected and non-infected plants with different phenotypes to remove local effects and then perform controlled infections. In addition, a wide variety of soil biota come into contact with wounded roots. In fact, we found entomopathogenic nematodes surrounding galls, which are likely to prey on weevil remains but may also affect weevil-plant interactions. Thus, identifying the relative effect of each interacting species on the complex network of hormones driving host response and phenotypic change is challenging.

Regardless of the mechanism underlying phenotypic changes, we found marked inter-individual variation in an *E. repandum* population where an unexpected pollination landscape has emerged. The lack of genetic exchange among non-infected plants, which maintain the phenotype associated with the selfing syndrome, contrasts with the pollinator movements among infected plants with large flowers, potentially leading to the accumulation of genetic differences among these two groups of individuals within the same population. In our study system, the increase in pollinator visitation is an indirect effect of the phenotype change that could result in an advantage for offspring production. An indirect non-negative effect on host functionality derived from weevil infection has also been described in infected individuals of *Cuscuta campestris*, where photosynthetic activity is increased. We hypothesize that pollinators would act as drivers of ecological divergence among sympatric individuals, in an analogous manner to the parasite-driven wedge model in animals, as we found evidence of assortative mating in this population.

We also hypothesize a shift from an antagonistic interaction mediated by weevil infection towards a mutualism assuming that pollination would benefit plant success. Recently, macroevolutionary studies carried out in Curculioninae subfamily of weevils support the benefit for the plant for having weevils larvae feeding in plant tissues (Haran et al. 2023). Brood-site (or nursery) pollination is a peculiar mutualism in which the plant provides a brood-site to the insect as a reward for being pollinated. This intimate interaction begins early in the life cycle of the insect because the larva develops directly in the host plant tissue. At an ancestral stage, Curculioninae weevil larvae fed on reproductive organs of the plant, but there are transitions to develop in root or stems.

Likewise, we provide new data on the cascading connection among species and highlight the importance of studying the species networks established both below and above ground to obtain a more realistic view of ecological evolutionary contexts in plants and a greater understanding of mechanisms promoting and maintaining species diversity. Since the loss of interactions among species also accompanies their extinction, an exhaustive understanding of species interaction dynamics is paramount in the current context of biodiversity emergency.

## Conclusions

In this work, we demonstrated pronounced inter-individual variation and its effect in modulating interactions with flower visitors, which potentially play an important role in offspring production in plants. Although weevil infection was not proven to be the cause of such variation, we demonstrated the occurrence of multiple interactions established by the same host plant. A differential pollination activity between infected and non-infected plants— or at least between plants showing phenotypic changes compared to the original phenotype—modulates gene flow within the population. Thus, the genetic structure of this population with known phenotypic variation is expected to differ from a population formed by the original phenotype of *E. repandum*, which is predominantly self-pollinated. Therefore, evolutionary trajectories might differ between groups with distinct phenotypes, leading to a substantial change in the dominant mating system of the species. Following the hypothesis of phenotypic change caused by weevil infection in this study, we illustrated how an *a priori* antagonistic interaction could positively influence the net performance of the host through its effect on subsequent species interactions. It highlights the dynamic nature of species interactions when they are part of multidimensional networks of relationships in wild communities.

**Figure 8.**
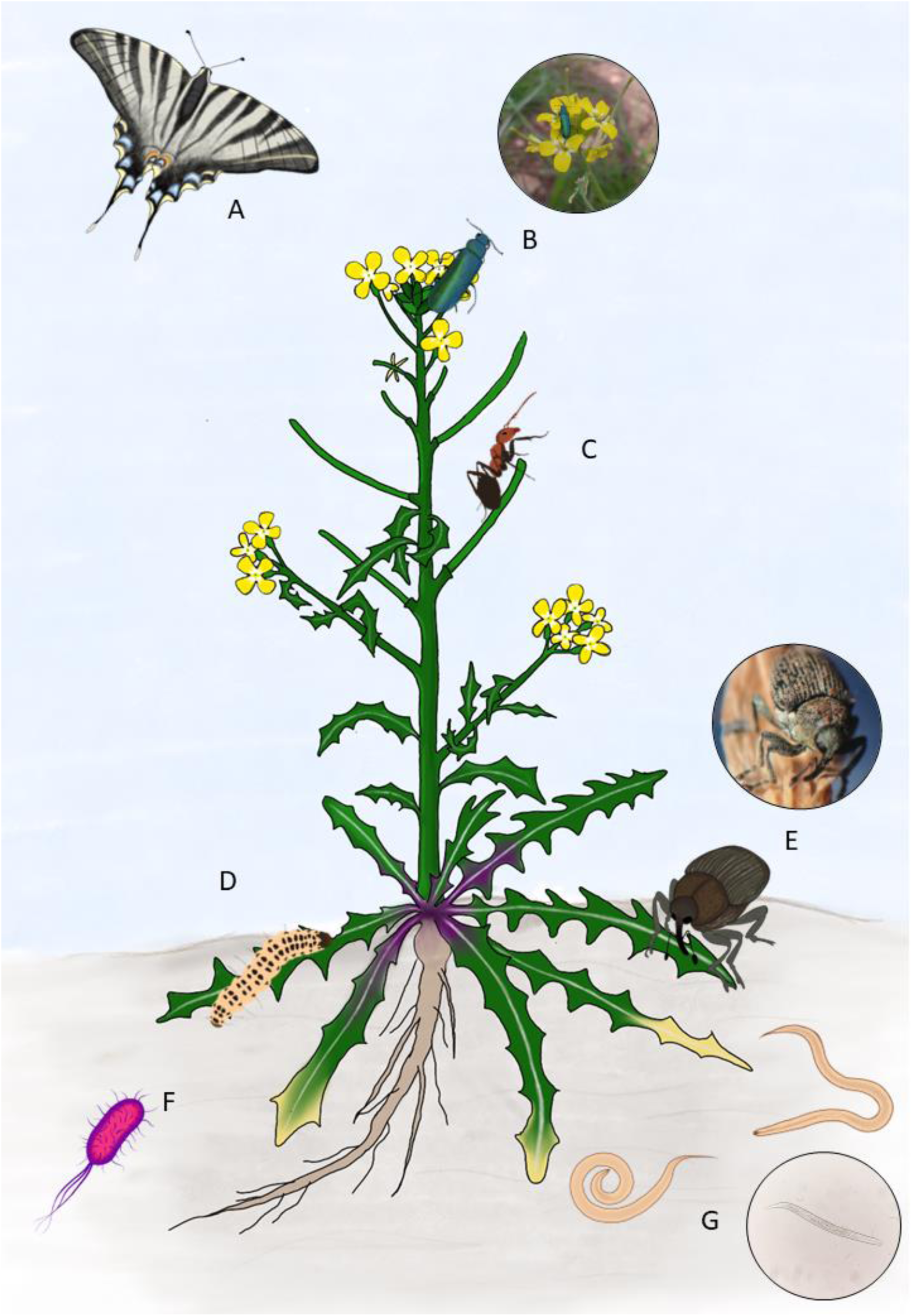
Picture illustrating some examples of below- and above-ground interactions potentially established by a host plant: (A) mutualism mediated by pollinators, (B) antagonism mediated by florivores, (C) mutualism mediated by organisms involved in defense against host herbivores, (D) antagonism mediated by herbivores, and (E-F-G) a wide variety of interactions from mutualism to antagonism mediated by root infection.

## References

Aboubakar Souna D, Bokonon-Ganta AH, Dannon EA, et al. 2019. Volatiles from Maruca vitrata (Lepidoptera, Crambidae) host plants influence olfactory responses of the parasitoid Therophilus javanus (Hymenoptera, Braconidae, Agathidinae). Biological control: theory and applications in pest management 130: 104–109.

Acosta IF, Gasperini D, Chételat A, Stolz S, Santuari L, Farmer EE. 2013. Role of NINJA in root jasmonate signaling. Proceedings of the National Academy of Sciences of the United States of America 110: 15473–15478.

Ali JG, Agrawal AA. 2012. Specialist versus generalist insect herbivores and plant defense. Trends in plant science 17: 293–302.

Altshuler DL. 1999. Novel interactions of non-pollinating ants with pollinators and fruit consumers in a tropical forest. Oecologia 119: 600–606.

Andriuzzi WS, Schmidt O, Brussaard L, Faber JH, Bolger T. 2016. Earthworm functional traits and interspecific interactions affect plant nitrogen acquisition and primary production. Applied soil ecology: a section of Agriculture, Ecosystems & Environment 104: 148–156.

Ansari MA, Shah FA, Butt TM. 2008. Combined use of entomopathogenic nematodes andMetarhizium anisopliaeas a new approach for black vine weevil,Otiorhynchus sulcatus, control. Entomologia experimentalis et applicata 129: 340–347.

Arroyo-Correa B, Jordano P, Bartomeus I. 2023. Intraspecific variation in species interactions promotes the feasibility of mutualistic assemblages. Ecology letters 26: 448–459.

Ashman T-L, Majetic CJ. 2006. Genetic constraints on floral evolution: a review and evaluation of patterns. Heredity 96: 343–352.

Berendsen RL, Pieterse CMJ, Bakker PAHM. 2012. The rhizosphere microbiome and plant health. Trends in plant science 17: 478–486.

Blüthgen N, Menzel F, Blüthgen N. 2006. Measuring specialization in species interaction networks. BMC ecology 6: 9.

Bonfante P, Genre A. 2010. Mechanisms underlying beneficial plant–fungus interactions in mycorrhizal symbiosis. Nature communications 1: 1–11.

Bronstein JL. 2009. The evolution of facilitation and mutualism. The Journal of ecology 97: 1160–1170.

Bronstein JL. 2015. Mutualism. Oxford University Press.

Bruinsma M, Lucas-Barbosa D, ten Broeke CJM, et al. 2014. Folivory affects composition of nectar, floral odor and modifies pollinator behavior. Journal of chemical ecology 40: 39–49.

Caballero, A., 2020. Quantitative genetics. Cambridge University Press.

Cardel YJ, Koptur S. 2010. Effects of Florivory on the Pollination of Flowers: An Experimental Field Study with a Perennial Plant. International journal of plant sciences 171.

Chamberlain SA, Bronstein JL, Rudgers JA. 2014. How context dependent are species interactions? Ecology letters 17: 881–890.

Cozzolino S, Fineschi S, Litto M, Scopece G, Trunschke J, Schiestl FP. 2015. Herbivory Increases Fruit Set in Silene latifolia: A Consequence of Induced Pollinator-Attracting Floral Volatiles? Journal of chemical ecology 41: 622–630.

Danderson CA, Molano-Flores B. 2010. Effects of Herbivory and Inflorescence Size on Insect Visitation to Eryngium yuccifolium (Apiaceae) a Prairie Plant. The American midland naturalist 163: 234–246.

Dannon EA, Tamò M, Van Huis A, Dicke M. 2010. Effects of volatiles from Maruca vitrata larvae and caterpillar-infested flowers of their host plant Vigna unguiculata on the foraging behavior of the parasitoid Apanteles taragamae. Journal of chemical ecology 36: 1083–1091.

Detrey J, Cognard V, Djian-Caporalino C, et al. 2022. Growth and root-knot nematode infection of tomato are influenced by mycorrhizal fungi and earthworms in an intercropping cultivation system with leeks. Applied soil ecology: a section of Agriculture, Ecosystems & Environment 169: 104181.

Dieckmann, L. 1972. Beiträge zur Insektenfauna der DDR: Coleoptera - Curculionidae: Ceutorhynchinae. Beiträge zur Entomologie 22: 3–12

Domínguez Núñez JA, Serrano JS, Barreal JAR, González JAS de O. 2006. The influence of mycorrhization with Tuber melanosporum in the afforestation of a Mediterranean site with Quercus ilex and Quercus faginea. Forest ecology and management 231: 226– 233.

Ekesi S. 2000. Effect of volatiles and crude extracts of different plant materials on egg viability ofmaruca vitrata andclavigralla tomentosicollis. Phytoparasitica; Israel journal of plant protection sciences 28: 305–310.

Elzinga JA, Atlan A, Biere A, Gigord L, Weis AE, Bernasconi G. 2007. Time after time: flowering phenology and biotic interactions. Trends in ecology & evolution 22: 432–439.

Feliner GN. 1991. Breeding systems and related floral traits in several Erysimum (Cruciferae). Canadian journal of botany. Journal canadien de botanique 69: 2515–2521.

Felten, J., Kohler, A., Morin, E., Bhalerao, R.P., Palme, K., Martin, F., Ditengou, F.A. and Legué, V., 2009. The ectomycorrhizal fungus Laccaria bicolor stimulates lateral root formation in poplar and *Arabidopsis* through auxin transport and signaling. Plant physiology, 151(4), pp.1991–2005.

Fernandez C, Santonja M, Gros R, et al. 2013. Allelochemicals of Pinus halepensis as drivers of biodiversity in Mediterranean open mosaic habitats during the colonization stage of secondary succession. Journal of chemical ecology 39: 298–311.

Fordyce, J.A., 2006. The evolutionary consequences of ecological interactions mediated through phenotypic plasticity. Journal of Experimental Biology, 209(12), pp.2377–2383.

García-Muñoz, A., Ferrón, C., Vaca-Benito, C., Loureiro, J., Castro, S., Muñoz-Pajares, A.J. and Abdelaziz, M., 2023. Ploidy effects on the relationship between floral phenotype, reproductive investment, and fitness in an autogamous species complex. American Journal of Botany, 110(6), p.e16197.

GBIF: The Global Biodiversity Information Facility (year) What is GBIF? Available from https://www.gbif.org/what-is-gbif [13 January 2020].

Gegear RJ, Manson JS, Thomson JD. 2007. Ecological context influences pollinator deterrence by alkaloids in floral nectar. Ecology letters 10: 375–382.

Giron D, Huguet E, Stone GN, Body M. 2016. Insect-induced effects on plants and possible effectors used by galling and leaf-mining insects to manipulate their host-plant. Journal of insect physiology 84: 70–89.

Gómez JM, Iriondo JM, Torres P. 2023. Modeling the continua in the outcomes of biotic interactions. Ecology 104: e3995.

Grant V. 1949. Pollination systems as isolating mechanisms in angiosperms. Evolution; international journal of organic evolution 3: 82–97.

Guerra-Guimarães L, Pinheiro C, Oliveira ASF, et al. 2023. The chloroplast protein HCF164 is predicted to be associated with Coffea SH9 resistance factor against Hemileia vastatrix. Scientific reports 13: 16019.

Haran J, Li X, Allio R, et al. 2023. Phylogenomics illuminates the phylogeny of flower weevils (Curculioninae) and reveals ten independent origins of brood-site pollination mutualism in true weevils. Proceedings. Biological sciences / The Royal Society 290: 20230889.

Hardoim, P.R., van Overbeek, L.S. and van Elsas, J.D., 2008. Properties of bacterial endophytes and their proposed role in plant growth. Trends in microbiology, 16(10), pp.463–471.

Heil M, Fiala B, Baumann B, Linsenmair KE. 2000. Temporal, Spatial and Biotic Variations in Extrafloral Nectar Secretion by *Macaranga tanarius*. Functional ecology 14: 749–757.

Heil M, Koch T, Hilpert A, Fiala B, Boland W, Linsenmair K. 2001. Extrafloral nectar production of the ant-associated plant, Macaranga tanarius, is an induced, indirect, defensive response elicited by jasmonic acid. Proceedings of the National Academy of Sciences of the United States of America 98: 1083–1088.

Herrera CM, Pellmyr O. 2009. Plant Animal Interactions: An Evolutionary Approach. John Wiley & Sons.

Hogenhout, S.A. and Bos, J.I., 2011. Effector proteins that modulate plant–insect interactions. Current opinion in plant biology, 14(4), pp.422–428.

Johnson SN, Erb M, Hartley SE. 2016. Roots under attack: contrasting plant responses to below- and aboveground insect herbivory. The New phytologist 210: 413–418.

Jordano P. 2016a. Sampling networks of ecological interactions. Functional ecology 30: 1883– 1893.

Jordano P. 2016b. Chasing Ecological Interactions. PLoS biology 14: e1002559.

Kaczorowski RL, Juenger TE, Holtsford TP. 2008. Heritability and correlation structure of nectar and floral morphology traits in Nicotiana alata. Evolution; international journal of organic evolution 62: 1738–1750.

Kammerhofer N, Radakovic Z, Regis JMA, et al. 2015. Role of stress-related hormones in plant defence during early infection of the cyst nematode Heterodera schachtii in Arabidopsis. The New phytologist 207: 778–789.

Leach JE, Triplett LR, Argueso CT, Trivedi P. 2017. Communication in the Phytobiome. Cell 169: 587–596.

Lugtenberg B, Kamilova F. 2009. Plant-growth-promoting rhizobacteria. Annual review of microbiology 63: 541–556.

Lu J, Robert CAM, Riemann M, et al. 2015. Induced jasmonate signaling leads to contrasting effects on root damage and herbivore performance. Plant physiology 167: 1100–1116.

Mandyam KG, Jumpponen A. 2014. Mutualism-parasitism paradigm synthesized from results of root-endophyte models. Frontiers in microbiology 5: 776.

Manzaneda AJ, Rey PJ. 2009. Assessing ecological specialization of an ant-seed dispersal mutualism through a wide geographic range. Ecology 90: 3009–3022.

McNaughton SJ. 1983. Compensatory plant growth as a response to herbivory. Oikos 40: 329.

Meier AR, Hunter MD. 2018. Arbuscular mycorrhizal fungi mediate herbivore-induction of plant defenses differently above and belowground. Oikos 127: 1759–1775.

Minnaar C, de Jager ML, Anderson B. 2019. Intraspecific divergence in floral-tube length promotes asymmetric pollen movement and reproductive isolation. The New phytologist 224: 1160–1170.

Morales FJ, Tamayo PJ, Castaño M, Olaya C, Martínez AK, Velasco AC. 2009. Enfermedades virales del tomate (Solanum Lycopersicum L.) en Colombia. Fitopatologia colombiana: Revista de la Asociacion Colombiana de Fitopalogia y Ciencias Afines “ASCOLFI.” 33: 23–27.

Moran NP, Caspers BA, Chakarov N, et al. 2022. Shifts between cooperation and antagonism driven by individual variation: a systematic synthesis review. Oikos 2022.

Moreira X, Abdala-Roberts L, Parra-Tabla V, Mooney KA. 2015. Latitudinal variation in herbivory: influences of climatic drivers, herbivore identity and natural enemies. Oikos 124: 1444–1452.

Moreira X, Castagneyrol B, Abdala-Roberts L, Traveset A. 2019. A meta-analysis of herbivore effects on plant attractiveness to pollinators. Ecology 100: e02707.

Muchhala N, Potts MD. 2007. Character displacement among bat-pollinated flowers of the genus Burmeistera: analysis of mechanism, process and pattern. Proceedings. Biological sciences / The Royal Society 274: 2731–2737.

Muñoz-Gallego R, Fedriani JM, Serra PE, Traveset A. 2022. Nonadditive effects of two contrasting introduced herbivores on the reproduction of a pollination-specialized palm. Ecology 103: e3797.

Murakami R, Ushima R, Sugimoto R, et al. 2021. A new galling insect model enhances photosynthetic activity in an obligate holoparasitic plant. Scientific reports 11: 13013.

Nakazawa T. 2017. Individual interaction data are required in community ecology: a conceptual review of the predator–prey mass ratio and more. Ecological research 32: 5–12.

Ness JH. 2006. A mutualism’s indirect costs: the most aggressive plant bodyguards also deter pollinators. Oikos 113: 506–514.

Newingham BA, Callaway RM, Bassirirad H. 2007. Allocating nitrogen away from a herbivore: a novel compensatory response to root herbivory. Oecologia 153: 913–920.

Oberprieler RG, Marvaldi AE, Anderson RS. 2007. Weevils, weevils, weevils everywhere. Zootaxa 1668: 491–520.

Opedal, Ø.H., Armbruster, W.S., Hansen, T.F., Holstad, A., Pélabon, C., Andersson, S., Campbell, D.R., Caruso, C.M., Delph, L.F., Eckert, C.G. and Lankinen, Å., 2023. Evolvability and trait function predict phenotypic divergence of plant populations. Proceedings of the National Academy of Sciences, 120(1), p.e2203228120.

Perea R, Delibes M, Polko M, Suárez-Esteban A, Fedriani JM. 2013. Context-dependent fruit-frugivore interactions: partner identities and spatio-temporal variations. Oikos 122: 943–951.

Pilosof S, Porter MA, Pascual M, Kéfi S. 2017. The multilayer nature of ecological networks. Nature ecology & evolution 1: 101.

Poisot T, Stanko M, Miklisová D, Morand S. 2013. Facultative and obligate parasite communities exhibit different network properties. Parasitology 140: 1340–1345.

Poveda K, Steffan-Dewenter I, Scheu S, Tscharntke T. 2003. Effects of below- and above-ground herbivores on plant growth, flower visitation and seed set. Oecologia 135: 601–605.

Preston DL. 2010. Ecological consequences of parasitism. http://johnsonlaboratory.com/sites/default/files/publications/Preston%20and%20Johnson%202010.pdf. 14 Sep. 2023.

Price PW, Westoby M, Rice B, et al. 1986. Parasite mediation in ecological interations. Annual review of ecology and systematics 17: 487–505.

Quilbé J, Montiel J, Arrighi J-F, Stougaard J. 2022. Molecular Mechanisms of Intercellular Rhizobial Infection: Novel Findings of an Ancient Process. Frontiers in plant science 13: 922982.

R Core Team. 2023. R: A language and environment for statistical computing. R Foundation for Statistical. Computing, Vienna, Austria. URL https://www.R-project.org/.

Rasmann S, Bennett A, Biere A, Karley A, Guerrieri E. 2017. Root symbionts: Powerful drivers of plant above- and belowground indirect defenses. Insect science 24: 947–960.

Rasmann S, Turlings TC. 2016. Root signals that mediate mutualistic interactions in the rhizosphere. Current opinion in plant biology 32: 62–68.

Rose SJ, Burnside OC, Specht JE, Swisher BA. 1984. Competition and allelopathy between soybeans and weeds^1^. Agronomy journal 76: 523–528.

Rundle HD, Nosil P. 2005. Ecological speciation. Ecology letters 8: 336–352.

Rymer PD, Johnson SD, Savolainen V. 2010. Pollinator behaviour and plant speciation: can assortative mating and disruptive selection maintain distinct floral morphs in sympatry? The New phytologist 188: 426–436.

Saikkonen K, Faeth SH, Helander M, Sullivan TJ. 1998. FUNGAL ENDOPHYTES: A Continuum of Interactions with Host Plants. Annual review of ecology and systematics 29: 319–343.

Sapp J. 2004. The dynamics of symbiosis: an historical overview. Canadian journal of botany. Journal canadien de botanique 82: 1046–1056.

Sawaya GM, Goldberg AS, Steele MA, Dalgleish HJ. 2018. Environmental variation shifts the relationship between trees and scatterhoarders along the continuum from mutualism to antagonism. Integrative zoology 13: 319–330.

Scherf, H. 1964. Die Entwicklungsstadien der mitteleuropäischen Curculioniden (Morphologie, Bionomie, Ökologie). Abhandlungen der Senckenbergischen Naturforschenden Gesellschaft, 506: 1–335 pp.

Schulz B. 2006. Mutualistic interactions with fungal root endophytes. Microbial root endophytes: 261–279.

Seng S, Ponce GE, Andreas P, et al. 2023. Abscisic Acid: A Potential Secreted Effector Synthesized by Phytophagous Insects for Host-Plant Manipulation. Insects 14.

Silva M do C, Várzea V, Guerra-Guimarães L, et al. 2006. Coffee resistance to the main diseases: leaf rust and coffee berry disease. Revista brasileira de fisiologia vegetal 18: 119–147.

Song C, Von Ahn S, Rohr RP, Saavedra S. 2020. Towards a Probabilistic Understanding About the Context-Dependency of Species Interactions. Trends in ecology & evolution 35: 384–396.

Stotz GC, Salgado-Luarte C, Escobedo VM, Valladares F, Gianoli E. 2021. Global trends in phenotypic plasticity of plants. Ecology letters 24: 2267–2281.

Strauss SY, Rudgers JA, Lau JA, Irwin RE. 2002. Direct and ecological costs of resistance to herbivory. Trends in ecology & evolution 17: 278–285.

Strauss SY, Siemens DH, Decher MB, Mitchell-Olds T. 1999. Ecological costs of plant resistance to herbivores in the currency of pollination. Evolution; international journal of organic evolution 53: 1105–1113.

Sukno SA, García VM, Shaw BD, Thon MR. 2008. Root infection and systemic colonization of maize by Colletotrichum graminicola. Applied and environmental microbiology 74: 823–832.

Talekar NS. 1982. Effects of a sweetpotato weevil (Coleoptera: Curculionidae) infestation on sweet potato root yields. Journal of economic entomology 75: 1042–1044.

Thamer S, Schädler M, Bonte D, Ballhorn DJ. 2011. Dual benefit from a belowground symbiosis: nitrogen fixing rhizobia promote growth and defense against a specialist herbivore in a cyanogenic plant. Plant and soil 341: 209–219.

Thompson JN. 1999. The evolution of species interactions. Science 284: 2116–2118.

Thornhill R, Fincher CL. 2013. The parasite-driven-wedge model of parapatric speciation. Journal of zoology 291: 23–33.

Tiffin P. 2000. Mechanisms of tolerance to herbivore damage: what do we know? Evolutionary ecology.

Toby Kiers E, Adler LS, Grman EL, Van Der Heijden MGA. 2010. Manipulating the jasmonate response: How do methyl jasmonate additions mediate characteristics of aboveground and belowground mutualisms? Functional ecology 24: 434–443.

Tylianakis JM, Didham RK, Bascompte J, Wardle DA. 2008. Global change and species interactions in terrestrial ecosystems. Ecology letters 11: 1351–1363.

Vale WS do, Neto B de MS, Araújo LR, et al. 2023. Anthonomus grandis grandis Boheman (Coleoptera: Curculionidae) induces the formation of shelter structures in cotton plants. Research Square.

Valiente-Banuet A, Aizen MA, Alcántara JM, et al. 2015. Beyond species loss: the extinction of ecological interactions in a changing world. Functional ecology 29: 299–307.

Valverde J, Gómez JM, Perfectti F. 2016. The temporal dimension in individual-based plant pollination networks. Oikos 125: 468–479.

Van der Niet T, Peakall R, Johnson SD. 2014. Pollinator-driven ecological speciation in plants: new evidence and future perspectives. Annals of botany 113: 199–211.

Van der Putten WH, Vet LEM, Harvey JA, Wäckers FL. 2001. Linking above- and belowground multitrophic interactions of plants, herbivores, pathogens, and their antagonists. Trends in ecology & evolution 16: 547–554.

Vannette RL, Hunter MD. 2009. Mycorrhizal fungi as mediators of defence against insect pests in agricultural systems. Agricultural and forest entomology 11: 351–358.

Vannette RL, Hunter MD. 2013. Mycorrhizal abundance affects the expression of plant resistance traits and herbivore performance. The Journal of ecology 101: 1019–1029.

Vannette RL, Hunter MD, Rasmann S. 2013. Arbuscular mycorrhizal fungi alter above- and below-ground chemical defense expression differentially among Asclepias species. Frontiers in plant science 4: 361.

Wang BC, Smith TB. 2002. Closing the seed dispersal loop. Trends in ecology & evolution 17: 379–386.

Zhang B, Sun S-F, Luo W-L, et al. 2021. A new brood-pollination mutualism between Stellera chamaejasme and flower thrips Frankliniella intonsa. BMC plant biology 21: 562.

Ponisio, L. C., Valdovinos, F. S., Allhoff, K. T., Gaiarsa, M. P., Barner, A., Guimarães Jr, P. R., … & Gillespie, R. (2019). A network perspective for community assembly. Frontiers in Ecology and Evolution, 7, 103.

Waser NM, Ollerton J, Eds: Plant-pollinator interactions: from specialization to generalization. 2006, Chicago: University of Chicago Press

Yeakel, J. D., Pires, M. M., de Aguiar, M. A., O’Donnell, J. L., Guimarães Jr, P. R., Gravel, D., & Gross, T. (2020). Diverse interactions and ecosystem engineering can stabilize community assembly. Nature communications, 11(1), 3307.

